## Supplementary Material for "A High-Quality De *Novo* Genome Assembly from a Single Mosquito using PacBio Sequencing"

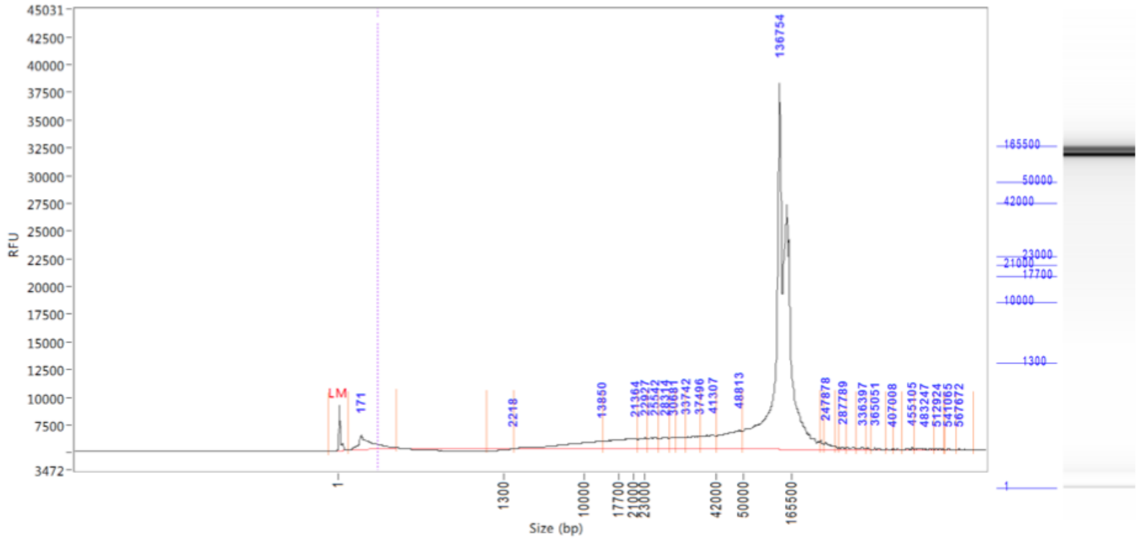


Figure S1. Femto Pulse evaluation of the Modified MagAttract DNA extraction prior to shipment to California.

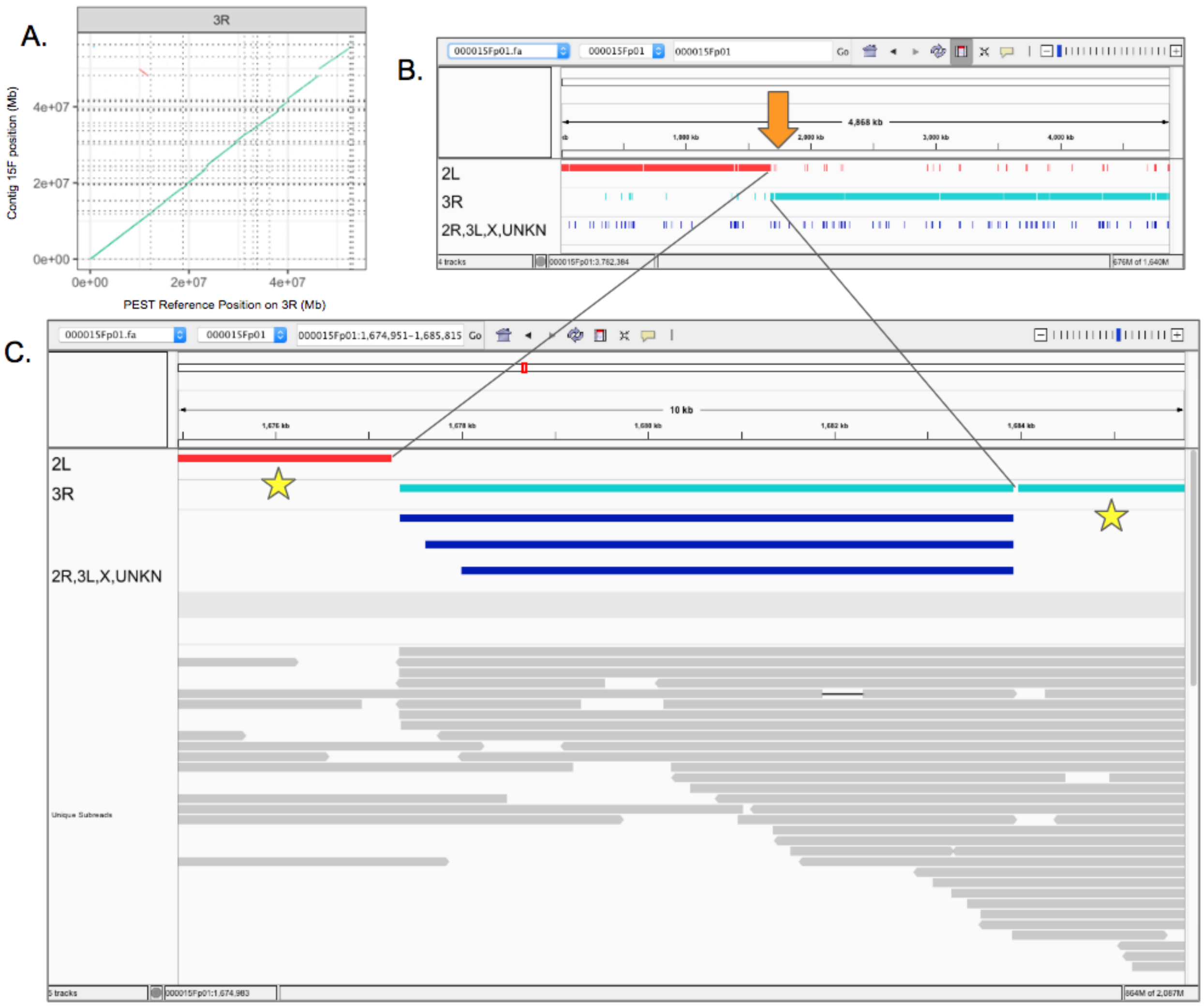


Figure S2. A chimeric contig between 2L and 3R. A. Alignment of PacBio contigs to PEST identifies a candidate chromosomal rearrangement. B. IGV screenshot of breakpoint (orange arrow) localized by alignment of contig to PEST. Red: alignment to 2L, turquoise: alignments to 3R, navy blue: alignments to other chromosomes and unplaced contigs. C. IGV visualization of mapped unique subreads at breakpoint shows 0 subreads mapping across the central repetitive region into the unique flanking sequence on the left (2L) and right (3R) (stars). A count of spanning reads was also determined with bedtools bamtobed utility. The 6.5kb central region aligns to four loci in the PEST genome and has ~370 bp of sequence similarity to the Tc1-like transposase gene in *An. gambiae*.

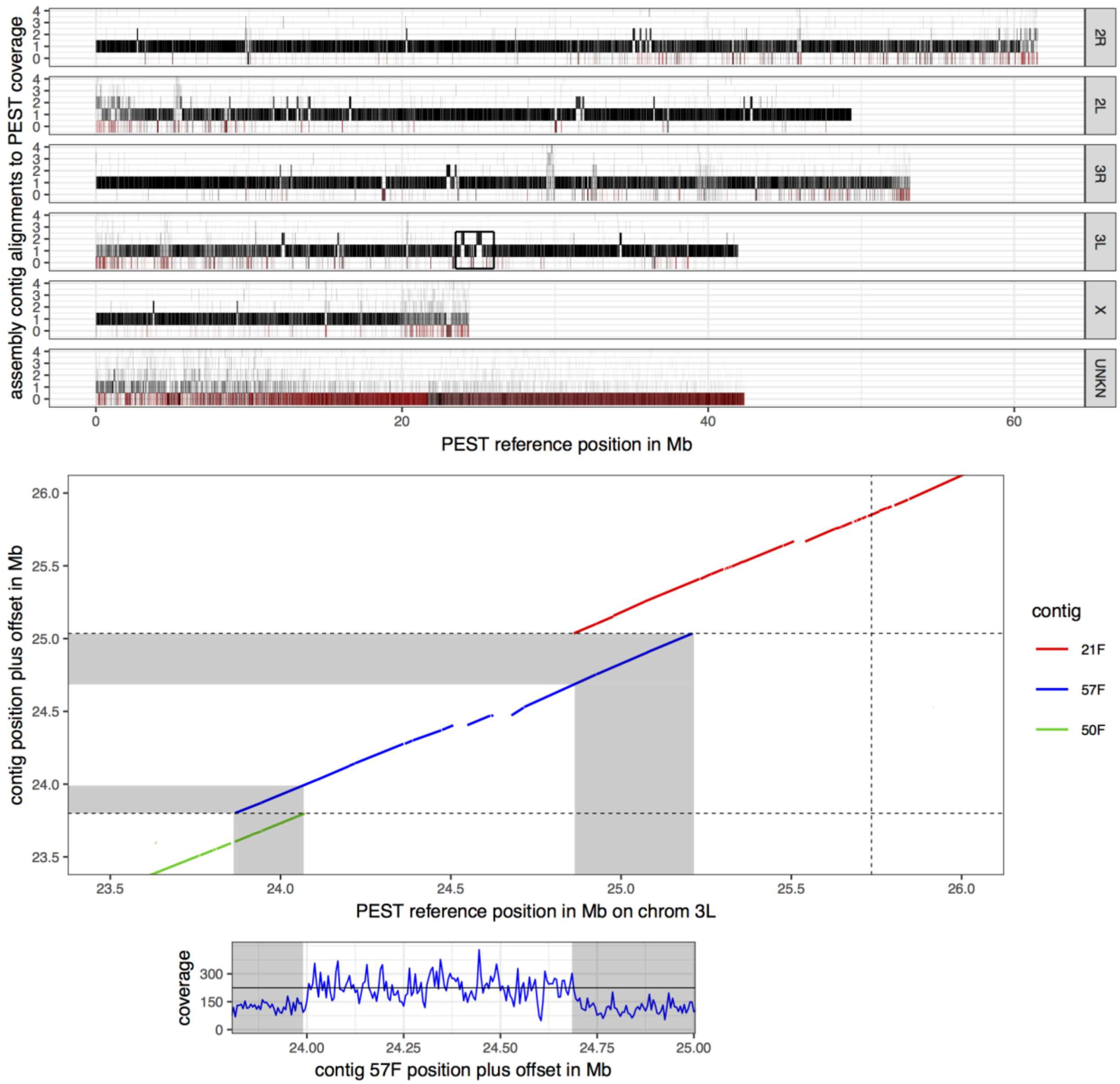


Figure S3. Alignment and coverage plot (top) of the PacBio assembly contigs relative to PEST, and magnification of one area of excess coverage (bottom). In the top panel, the number of alignments of PacBio contigs to PEST are represented by black bars, with most of the genome showing a 1:1 correspondence to PEST. Red denotes N's in the reference. Isolated areas of higher number of contig alignments are visible, one of which (black box) is magnified in the bottom panel. Here, the ends of neighboring contigs overlap, which is currently not resolved with the Purge Haplotigs software since the overlap is only partial. The sequencing depth of PacBio reads for the central (blue) contig (57F) corroborate this interpretation, exhibiting half of the expected coverage in the greyed regions of contig overlap, and with the corresponding ends of the red and green contigs complementing with the other half of coverage, respectively (not shown for clarity).

| | 1 Cell | 2 Cells | 3 Cells |
|---|---|---|---|
| **Total bases (Gb)** | 23.6 | 48.5 | 72.7 |
| **Total unique bases (Gb)** | 4.46 | 8.31 | 12.8 |
| **Unique coverage** | 17 X | 31 X | 45 X |
| **Assembly size (Mb)** | 150 | 265 | 271 |
| **Number of contigs** | 3,290 | 815 | 580 |
| **Contig N50 (Mb)** | 0.066 | 1.5 | 3.5 |

Table S1. Statistics for *Anopheles coluzzii de novo* genome assemblies as a function of the number of SMRT Cells used for the assembly.

| Loading concentration | Gb/ Cell | Mean Polymerase Read Length | N50 Polymerase Read Length | Mean Subread Length | N50 Subread Length | P0 | P1 | P2 |
|---|---|---|---|---|---|---|---|---|
| 5 pM | 24.1 | 40290 | 116615 | 8185 | 12978 | 26.0% | 60.1% | 13.9% |

| | | | | | | | | |
|---|---|---|---|---|---|---|---|---|
| 5 pM | 23.6 | 40077 | 114807 | 8254 | 13132 | 27.1% | 59.0% | 14.0% |
| 6 pM | 25.0 | 47177 | 122898 | 8012 | 12751 | 35.3% | 53.1% | 11.7% |

Table S2. Run statistics for the three Sequel SMRT Cells.

| **Gene Count (%)** | **Raw PacBio** | **Curated PacBio** | **Previous Reference** |
|---|---|---|---|
| Complete | 2745 (98.0) | 2747 (98.1) | 2448 (87.5) |
| Complete Single Copy | 2635 (94.1) | 2680 (95.7) | 2446 (87.4) |
| Complete Duplicated | 110 (3.9) | 67 (2.4) | 3 (0.1) |
| Fragmented | 25 (0.9) | 25 (0.9) | 190 (6.8) |
| Missing | 29 (1.1) | 28 (1.0) | 160 (5.7) |
| Total | 2799 (100) | 2799 (100) | 2799 (100) |

Table S3. Analysis of single copy conserved genes using BUSCO v3.0.2 and the diptera gene set. Raw PacBio: primary contigs from the 3-cell *de novo* FALCON-Unzip assembly. Curated PacBio: Primary contigs after removal of bacterial contaminants and duplicated haplotypes. Previous reference from [18] GCA_000150765.1.

Table S4: Over 90% of genes formerly unassigned to chromosomes (e.g., residing on the UNKN chromosome) have now been placed to chromosomes and contigs. This table shows the contig that each UNKN gene is found in, as well as the chromosome that the gene is now assigned to.

| gene | chrom | contig |
|---|---|---|
| AGAP012735 | 3R | 000000F |
| AGAP012478 | 3R | 000000F |
| AGAP012477 | 3R | 000000F |
| AGAP012476 | 3R | 000000F |
| AGAP012475 | 3R | 000000F |
| AGAP012474 | 3R | 000000F |
| AGAP012585 | 3R | 000000F |
| AGAP012855 | 3R | 000000F |
| AGAP012637 | 3R | 000000F |
| AGAP012905 | 3R | 000000F |
| AGAP012724 | 3R | 000000F |
| AGAP012723 | 3R | 000000F |
| AGAP012553 | 3R | 000000F |
| AGAP012512 | 3R | 000000F |

| | | |
|---|---|---|
| AGAP012828 | 3R | 000000F |
| AGAP012829 | 3R | 000000F |
| AGAP012800 | 3R | 000000F |
| AGAP012457 | 3R | 000000F |
| AGAP012842 | 3R | 000000F |
| AGAP012754 | 3R | 000000F |
| AGAP012939 | 3R | 000000F |
| AGAP012542 | 2R | 000001F |
| AGAP012730 | 2R | 000001F |
| AGAP012497 | 2R | 000001F |
| AGAP012684 | 2R | 000001F |
| AGAP012537 | 2R | 000001F |
| AGAP012739 | 2R | 000002F |
| AGAP012563 | 2R | 000002F |
| AGAP012509 | 2R | 000002F |
| AGAP012508 | 2R | 000002F |
| AGAP013643 | 2R | 000002F |

| | | |
|---|---|---|
| AGAP012691 | 2R | 000002F |
| AGAP012698 | 2R | 000002F |
| AGAP012839 | 2R | 000002F |
| AGAP012838 | 2R | 000002F |
| AGAP012837 | 2R | 000002F |
| AGAP012836 | 2R | 000002F |
| AGAP012833 | 2R | 000002F |
| AGAP012819 | 2R | 000002F |
| AGAP012811 | 2R | 000002F |
| AGAP012872 | 2R | 000002F |
| AGAP028588 | 2R | 000002F |
| AGAP012634 | 2R | 000002F |
| AGAP012714 | 2R | 000002F |
| AGAP012783 | 2R | 000002F |
| AGAP012929 | 2R | 000002F |
| AGAP012925 | 2R | 000002F |
| AGAP012944 | 2R | 000002F |

| | | |
|---|---|---|
| AGAP012556 | 2R | 000002F |
| AGAP012703 | 2R | 000002F |
| AGAP012702 | 2R | 000002F |
| AGAP012575 | 2R | 000002F |
| AGAP012570 | 2R | 000002F |
| AGAP012761 | 2R | 000002F |
| AGAP012513 | 2R | 000002F |
| AGAP012514 | 2R | 000002F |
| AGAP012802 | 2R | 000002F |
| AGAP012809 | 2R | 000002F |
| AGAP012646 | 2R | 000002F |
| AGAP012643 | 2R | 000002F |
| AGAP012668 | 2R | 000002F |
| AGAP012860 | 2R | 000002F |
| AGAP012604 | 2R | 000002F |
| AGAP012605 | 2R | 000002F |
| AGAP012883 | 2R | 000002F |

| | | |
|---|---|---|
| AGAP012792 | 2R | 000002F |
| AGAP012914 | 2R | 000002F |
| AGAP012936 | 2R | 000002F |
| AGAP012732 | 2R | 000003F |
| AGAP012774 | 2R | 000003F |
| AGAP012818 | 2R | 000003F |
| AGAP012899 | 2R | 000003F |
| AGAP012898 | 2R | 000003F |
| AGAP012781 | 2R | 000003F |
| AGAP012906 | 2R | 000003F |
| AGAP012920 | 2R | 000003F |
| AGAP012824 | 2R | 000003F |
| AGAP012932 | 2R | 000003F |
| AGAP012505 | 3L | 000005F |
| AGAP012504 | 3L | 000005F |
| AGAP012813 | 3L | 000005F |
| AGAP012817 | 3L | 000005F |

| | | |
|---|---|---|
| AGAP012871 | 3L | 000005F |
| AGAP028580 | 3L | 000005F |
| AGAP028585 | 3L | 000005F |
| AGAP012676 | 3L | 000005F |
| AGAP012587 | 3L | 000005F |
| AGAP012850 | 3L | 000005F |
| AGAP012853 | 3L | 000005F |
| AGAP012892 | 3L | 000005F |
| AGAP012895 | 3L | 000005F |
| AGAP012894 | 3L | 000005F |
| AGAP012907 | 3L | 000005F |
| AGAP012729 | 3L | 000005F |
| AGAP012554 | 3L | 000005F |
| AGAP012555 | 3L | 000005F |
| AGAP012777 | 3L | 000005F |
| AGAP012686 | 3L | 000005F |
| AGAP012827 | 3L | 000005F |

| | | |
|---|---|---|
| AGAP012879 | 3L | 000005F |
| AGAP012749 | 3L | 000005F |
| AGAP012806 | 3L | 000005F |
| AGAP012867 | 3L | 000005F |
| AGAP012843 | 3L | 000005F |
| AGAP012887 | 3L | 000005F |
| AGAP012911 | 3L | 000005F |
| AGAP012913 | 3L | 000005F |
| AGAP012933 | 3L | 000005F |
| AGAP012544 | 2R | 000006F |
| AGAP012618 | 2R | 000006F |
| AGAP012737 | 3R | 000007F |
| AGAP012814 | 3R | 000007F |
| AGAP012679 | 3R | 000007F |
| AGAP012635 | 3R | 000007F |
| AGAP012728 | 3R | 000007F |
| AGAP012578 | 3R | 000007F |

| | | |
|---|---|---|
| AGAP012600 | 3R | 000007F |
| AGAP012731 | 2R | 000008F |
| AGAP012694 | 2R | 000008F |
| AGAP012785 | 2R | 000008F |
| AGAP012908 | 2R | 000008F |
| AGAP012683 | 2R | 000008F |
| AGAP012689 | 2R | 000008F |
| AGAP012803 | 2R | 000008F |
| AGAP012520 | 2R | 000009F |
| AGAP012771 | 2L | 000010F |
| AGAP012659 | 2L | 000010F |
| AGAP012658 | 2L | 000010F |
| AGAP028581 | 2L | 000010F |
| AGAP012677 | 2L | 000010F |
| AGAP012589 | 2L | 000010F |
| AGAP012580 | 2L | 000010F |
| AGAP012847 | 2L | 000010F |

| | | |
|---|---|---|
| AGAP012821 | 2L | 000012F |
| AGAP012547 | 2R | 000013F |
| AGAP012560 | 2R | 000013F |
| AGAP012718 | 2R | 000013F |
| AGAP012711 | 2R | 000013F |
| AGAP012586 | 2R | 000013F |
| AGAP012613 | 2R | 000013F |
| AGAP012710 | 2R | 000013F |
| AGAP012926 | 2R | 000013F |
| AGAP012558 | 2R | 000013F |
| AGAP012559 | 2R | 000013F |
| AGAP012705 | 2R | 000013F |
| AGAP012709 | 2R | 000013F |
| AGAP012574 | 2R | 000013F |
| AGAP012572 | 2R | 000013F |
| AGAP012573 | 2R | 000013F |
| AGAP012740 | 2R | 000013F |

| | | |
|---|---|---|
| AGAP012748 | 2R | 000013F |
| AGAP012841 | 2R | 000013F |
| AGAP012609 | 2R | 000013F |
| AGAP012882 | 2R | 000013F |
| AGAP012793 | 2R | 000013F |
| AGAP012930 | 2R | 000013F |
| AGAP012549 | 2R | 000014F |
| AGAP012561 | 2R | 000014F |
| AGAP012870 | 2R | 000014F |
| AGAP012877 | 2R | 000014F |
| AGAP012857 | 2R | 000014F |
| AGAP012868 | 2R | 000014F |
| AGAP012869 | 2R | 000014F |
| AGAP012611 | 2R | 000014F |
| AGAP012787 | 2R | 000014F |
| AGAP012922 | 2R | 000014F |
| AGAP012550 | 2R | 000014F |

| | | |
|---|---|---|
| AGAP012823 | 2R | 000014F |
| AGAP012746 | 2R | 000014F |
| AGAP012801 | 2R | 000014F |
| AGAP012641 | 2R | 000014F |
| AGAP012642 | 2R | 000014F |
| AGAP012866 | 2R | 000014F |
| AGAP012661 | 2R | 000014F |
| AGAP012662 | 2R | 000014F |
| AGAP012599 | 2R | 000014F |
| AGAP012590 | 2R | 000014F |
| AGAP012591 | 2R | 000014F |
| AGAP012750 | 2R | 000014F |
| AGAP012795 | 2R | 000014F |
| AGAP012799 | 2R | 000014F |
| AGAP012917 | 2R | 000014F |
| AGAP012712 | 3R | 000015F |
| AGAP012657 | 3R | 000015F |

| | | |
|---|---|---|
| AGAP012875 | 3R | 000015F |
| AGAP012644 | 3R | 000015F |
| AGAP012582 | 3R | 000015F |
| AGAP012581 | 3R | 000015F |
| AGAP012927 | 3R | 000015F |
| AGAP012606 | 3R | 000015F |
| AGAP012747 | 3R | 000015F |
| AGAP012743 | 3R | 000015F |
| AGAP012632 | 3R | 000015F |
| AGAP012669 | 3R | 000015F |
| AGAP012849 | 3R | 000015F |
| AGAP012628 | 3R | 000015F |
| AGAP012798 | 3R | 000015F |
| AGAP012564 | X | 000016F |
| AGAP012675 | X | 000016F |
| AGAP012715 | X | 000016F |
| AGAP012511 | X | 000016F |

| | | |
|---|---|---|
| AGAP012536 | X | 000016F |
| AGAP012535 | X | 000016F |
| AGAP012450 | X | 000016F |
| AGAP012888 | X | 000016F |
| AGAP012720 | 2R | 000018F |
| AGAP012805 | 2R | 000018F |
| AGAP012596 | 2R | 000018F |
| AGAP012645 | 3R | 000019F |
| AGAP012854 | 3L | 000021F |
| AGAP012893 | 3L | 000021F |
| AGAP012858 | 3L | 000021F |
| AGAP012943 | 3L | 000021F |
| AGAP012845 | 3L | 000021F |
| AGAP012881 | 3L | 000021F |
| AGAP028578 | X | 000022F |
| AGAP012545 | 3R | 000023F |
| AGAP012796 | 3R | 000023F |

| | | |
|---|---|---|
| AGAP012758 | 2R | 000025F |
| AGAP012812 | 2R | 000025F |
| AGAP028586 | 2R | 000025F |
| AGAP012859 | 2R | 000025F |
| AGAP012856 | 2R | 000025F |
| AGAP012713 | 2R | 000025F |
| AGAP012901 | 2R | 000025F |
| AGAP012928 | 2R | 000025F |
| AGAP012726 | 2R | 000025F |
| AGAP012571 | 2R | 000025F |
| AGAP012769 | 2R | 000025F |
| AGAP012820 | 2R | 000025F |
| AGAP012598 | 2R | 000025F |
| AGAP012620 | 2R | 000025F |
| AGAP012938 | 2R | 000025F |
| AGAP012501 | 3L | 000027F |
| AGAP012773 | 3L | 000027F |

| | | |
|---|---|---|
| AGAP012489 | 3L | 000027F |
| AGAP012863 | 3L | 000027F |
| AGAP012593 | 3L | 000027F |
| AGAP012456 | 3L | 000028F |
| AGAP013599 | 2L | 000029F |
| AGAP012695 | 2L | 000029F |
| AGAP012699 | 2L | 000029F |
| AGAP012529 | 2L | 000029F |
| AGAP012528 | 2L | 000029F |
| AGAP012775 | 2L | 000029F |
| AGAP012685 | 2L | 000029F |
| AGAP012530 | 2L | 000029F |
| AGAP012531 | 2L | 000029F |
| AGAP012804 | 2L | 000029F |
| AGAP012660 | 2L | 000029F |
| AGAP012759 | 2L | 000029F |
| AGAP012934 | 2L | 000029F |

| | | |
|---|---|---|
| AGAP012756 | 3R | 000030F |
| AGAP012690 | 3R | 000030F |
| AGAP012666 | 3R | 000030F |
| AGAP012434 | 3R | 000030F |
| AGAP012433 | 3R | 000030F |
| AGAP012437 | 3R | 000030F |
| AGAP012436 | 3R | 000030F |
| AGAP012435 | 3R | 000030F |
| AGAP028651 | 3R | 000030F |
| AGAP012603 | 3R | 000030F |
| AGAP012822 | 3R | 000030F |
| AGAP012636 | 3R | 000030F |
| AGAP012633 | 3R | 000030F |
| AGAP012755 | 3R | 000030F |
| AGAP012655 | 3R | 000031F |
| AGAP012673 | 3R | 000031F |
| AGAP012765 | 3R | 000031F |

| | | |
|---|---|---|
| AGAP012921 | 3R | 000031F |
| AGAP012649 | 3R | 000031F |
| AGAP012623 | 3R | 000031F |
| AGAP012915 | 3R | 000031F |
| AGAP012664 | 3L | 000032F |
| AGAP012770 | 3L | 000032F |
| AGAP012865 | 3L | 000032F |
| AGAP012640 | 3L | 000033F |
| AGAP012638 | 3L | 000033F |
| AGAP012579 | 3L | 000033F |
| AGAP012565 | 3L | 000034F |
| AGAP012566 | 3L | 000034F |
| AGAP012473 | 3L | 000034F |
| AGAP012472 | 3L | 000034F |
| AGAP012492 | 3L | 000034F |
| AGAP012471 | 3L | 000034F |
| AGAP012470 | 3L | 000034F |

| | | |
|---|---|---|
| AGAP012810 | 3L | 000034F |
| AGAP028583 | 3L | 000034F |
| AGAP028584 | 3L | 000034F |
| AGAP012469 | 3L | 000034F |
| AGAP012671 | 3L | 000034F |
| AGAP012670 | 3L | 000034F |
| AGAP012861 | 3L | 000034F |
| AGAP028704 | 3L | 000034F |
| AGAP028705 | 3L | 000034F |
| AGAP012780 | 3L | 000034F |
| AGAP012688 | 3L | 000034F |
| AGAP012626 | 3L | 000034F |
| AGAP012886 | 3L | 000034F |
| AGAP012797 | 3L | 000034F |
| AGAP012656 | 2L | 000035F |
| AGAP012525 | 2L | 000035F |
| AGAP012527 | 2L | 000035F |

| | | |
|---|---|---|
| AGAP012523 | 2L | 000035F |
| AGAP012496 | 2L | 000035F |
| AGAP012734 | 2L | 000035F |
| AGAP012612 | 2L | 000035F |
| AGAP012462 | 2L | 000035F |
| AGAP012763 | 2L | 000035F |
| AGAP012524 | 2L | 000035F |
| AGAP012707 | 2L | 000035F |
| AGAP012708 | 2L | 000035F |
| AGAP012495 | 2L | 000035F |
| AGAP012490 | 2L | 000035F |
| AGAP012493 | 2L | 000035F |
| AGAP012487 | 2L | 000035F |
| AGAP012534 | 2L | 000035F |
| AGAP012825 | 2L | 000035F |
| AGAP012826 | 2L | 000035F |
| AGAP012465 | 2L | 000035F |

| | | |
|---|---|---|
| AGAP012498 | 2L | 000035F |
| AGAP012486 | 2L | 000035F |
| AGAP028807 | 2L | 000036F |
| AGAP028810 | 2L | 000036F |
| AGAP012725 | 2L | 000037F |
| AGAP012653 | 3R | 000038F |
| AGAP012830 | 3R | 000038F |
| AGAP012674 | 3R | 000038F |
| AGAP012654 | 3R | 000038F |
| AGAP012706 | 3R | 000038F |
| AGAP012680 | 3R | 000038F |
| AGAP012681 | 3R | 000038F |
| AGAP012682 | 3R | 000038F |
| AGAP012807 | 3R | 000038F |
| AGAP012764 | 3R | 000038F |
| AGAP012736 | 2L | 000040F |
| AGAP012831 | 2L | 000040F |

| | | |
|---|---|---|
| AGAP012873 | 2L | 000040F |
| AGAP012569 | 2L | 000040F |
| AGAP012896 | 2L | 000040F |
| AGAP012903 | 2L | 000040F |
| AGAP012924 | 2L | 000040F |
| AGAP012667 | 2L | 000040F |
| AGAP012937 | 2L | 000040F |
| AGAP012562 | 3L | 000041F |
| AGAP012692 | 3L | 000041F |
| AGAP012716 | 3L | 000041F |
| AGAP012678 | 3L | 000041F |
| AGAP012576 | 3L | 000041F |
| AGAP012577 | 3L | 000041F |
| AGAP012900 | 3L | 000041F |
| AGAP012594 | 3L | 000041F |
| AGAP012627 | 3L | 000041F |
| AGAP012885 | 3L | 000041F |

| | | |
|---|---|---|
| AGAP012697 | 3L | 000042F |
| AGAP012696 | 3L | 000042F |
| AGAP012557 | 3L | 000042F |
| AGAP012851 | 3L | 000042F |
| AGAP012852 | 3L | 000042F |
| AGAP012789 | 3L | 000042F |
| AGAP012503 | 3L | 000042F |
| AGAP029093 | 3L | 000042F |
| AGAP012808 | 3L | 000042F |
| AGAP012648 | 3L | 000042F |
| AGAP012631 | 3L | 000042F |
| AGAP012844 | 3L | 000042F |
| AGAP012647 | 3L | 000042F |
| AGAP012880 | 3L | 000042F |
| AGAP012910 | 3L | 000042F |
| AGAP012639 | 3R | 000045F |
| AGAP012786 | 3R | 000045F |

| | | |
|---|---|---|
| AGAP012687 | 3R | 000045F |
| AGAP012693 | 3L | 000047F |
| AGAP012584 | 3L | 000047F |
| AGAP012788 | 3L | 000047F |
| AGAP012567 | 2L | 000049F |
| AGAP012651 | 2L | 000049F |
| AGAP012650 | 2L | 000049F |
| AGAP012616 | 2L | 000049F |
| AGAP012902 | 2L | 000049F |
| AGAP012941 | 2L | 000049F |
| AGAP012617 | 2L | 000049F |
| AGAP012779 | 2L | 000049F |
| AGAP012568 | 2L | 000049F |
| AGAP012546 | 3L | 000050F |
| AGAP012652 | 3L | 000050F |
| AGAP012834 | 3L | 000050F |
| AGAP012832 | 3L | 000050F |

| | | |
|---|---|---|
| AGAP012704 | 3L | 000050F |
| AGAP012816 | 3L | 000050F |
| AGAP028589 | 3L | 000050F |
| AGAP028587 | 3L | 000050F |
| AGAP012719 | 3L | 000050F |
| AGAP012772 | 3L | 000050F |
| AGAP012742 | 3L | 000050F |
| AGAP012753 | 3L | 000050F |
| AGAP012602 | 3L | 000050F |
| AGAP012741 | 3L | 000050F |
| AGAP012918 | 3L | 000050F |
| AGAP012919 | 3L | 000050F |
| AGAP012931 | 3L | 000050F |
| AGAP028799 | 3R | 000051F |
| AGAP028829 | X | 000052F |
| AGAP028828 | X | 000052F |
| AGAP028986 | X | 000052F |

| | | |
|---|---|---|
| AGAP028834 | X | 000052F |
| AGAP028990 | X | 000052F |
| AGAP028994 | X | 000052F |
| AGAP028859 | X | 000052F |
| AGAP028855 | X | 000052F |
| AGAP028856 | X | 000052F |
| AGAP028870 | X | 000052F |
| AGAP028837 | X | 000052F |
| AGAP028937 | X | 000052F |
| AGAP028936 | X | 000052F |
| AGAP028933 | X | 000052F |
| AGAP028919 | X | 000052F |
| AGAP028998 | X | 000052F |
| AGAP028956 | X | 000052F |
| AGAP028953 | X | 000052F |
| AGAP028952 | X | 000052F |
| AGAP028889 | X | 000052F |

| | | |
|---|---|---|
| AGAP028885 | X | 000052F |
| AGAP028985 | X | 000052F |
| AGAP028951 | X | 000052F |
| AGAP028846 | X | 000052F |
| AGAP028864 | X | 000052F |
| AGAP028861 | X | 000052F |
| AGAP028863 | X | 000052F |
| AGAP028929 | X | 000052F |
| AGAP028909 | X | 000052F |
| AGAP012453 | X | 000052F |
| AGAP028962 | X | 000052F |
| AGAP029035 | X | 000052F |
| AGAP028895 | X | 000052F |
| AGAP028896 | X | 000052F |
| AGAP028793 | X | 000052F |
| AGAP028797 | X | 000052F |
| AGAP028794 | X | 000052F |

| | | |
|---|---|---|
| AGAP028858 | X | 000052F |
| AGAP028912 | X | 000052F |
| AGAP028813 | X | 000052F |
| AGAP012722 | 3L | 000057F |
| AGAP012835 | 3L | 000057F |
| AGAP012619 | 3L | 000057F |
| AGAP012768 | 3L | 000057F |
| AGAP012608 | 3L | 000057F |
| AGAP012607 | 3L | 000057F |
| AGAP012767 | 3L | 000057F |
| AGAP028423 | 3L | 000057F |
| AGAP028826 | 2L | 000058F |
| AGAP028897 | 2L | 000058F |
| AGAP028839 | 2L | 000058F |
| AGAP028833 | 2L | 000058F |
| AGAP028993 | 2L | 000058F |
| AGAP028853 | 2L | 000058F |

| | | |
|---|---|---|
| AGAP028883 | 2L | 000058F |
| AGAP028948 | 2L | 000058F |
| AGAP028800 | 2L | 000058F |
| AGAP028821 | 2L | 000058F |
| AGAP028820 | 2L | 000058F |
| AGAP028823 | 2L | 000058F |
| AGAP028822 | 2L | 000058F |
| AGAP028825 | 2L | 000058F |
| AGAP028849 | 2L | 000058F |
| AGAP028848 | 2L | 000058F |
| AGAP028867 | 2L | 000058F |
| AGAP028862 | 2L | 000058F |
| AGAP028868 | 2L | 000058F |
| AGAP028924 | 2L | 000058F |
| AGAP028927 | 2L | 000058F |
| AGAP028922 | 2L | 000058F |
| AGAP028903 | 2L | 000058F |

| | | |
|---|---|---|
| AGAP028900 | 2L | 000058F |
| AGAP012451 | 2L | 000058F |
| AGAP028963 | 2L | 000058F |
| AGAP028943 | 2L | 000058F |
| AGAP028947 | 2L | 000058F |
| AGAP028944 | 2L | 000058F |
| AGAP029032 | 2L | 000058F |
| AGAP028892 | 2L | 000058F |
| AGAP028811 | 2L | 000058F |
| AGAP028792 | 2L | 000058F |
| AGAP028790 | 2L | 000058F |
| AGAP028928 | 2L | 000058F |
| AGAP028814 | 2L | 000058F |
| AGAP028815 | 2L | 000058F |
| AGAP028816 | 2L | 000058F |
| AGAP028817 | 2L | 000058F |
| AGAP028818 | 2L | 000058F |

| | | |
|---|---|---|
| AGAP028819 | 2L | 000058F |
| AGAP012543 | 2R | 000059F |
| AGAP012541 | 2R | 000059F |
| AGAP012439 | 2R | 000059F |
| AGAP012518 | 2R | 000059F |
| AGAP012540 | 2R | 000059F |
| AGAP028996 | 2L | 000060F |
| AGAP028991 | 2L | 000060F |
| AGAP012428 | 2L | 000060F |
| AGAP028803 | 2L | 000060F |
| AGAP028842 | 2L | 000060F |
| AGAP012432 | 2L | 000060F |
| AGAP012430 | 2L | 000060F |
| AGAP012429 | 2L | 000060F |
| AGAP012431 | 2L | 000060F |
| AGAP012521 | 3R | 000062F |
| AGAP012522 | 3R | 000062F |

| | | |
|---|---|---|
| AGAP012464 | 3R | 000062F |
| AGAP012454 | 2R | 000063F |
| AGAP012447 | 2R | 000063F |
| AGAP028577 | 2R | 000063F |
| AGAP028576 | 2R | 000063F |
| AGAP028806 | 2L | 000066F |
| AGAP028805 | 2L | 000066F |
| AGAP028804 | 2L | 000066F |
| AGAP012458 | 3L | 000071F |
| AGAP012904 | 3L | 000071F |
| AGAP012721 | 2L | 000073F |
| AGAP012945 | 2L | 000073F |
| AGAP012595 | 2L | 000073F |
| AGAP028995 | 2R | 000074F |
| AGAP012552 | 2R | 000074F |
| AGAP012622 | 2L | 000081F |
| AGAP012459 | 2L | 000081F |

| | | |
|---|---|---|
| AGAP012757 | 2R | 000083F |
| AGAP012745 | 2R | 000083F |
| AGAP012665 | 2R | 000083F |
| AGAP012717 | 2L | 000085F |
| AGAP012519 | 2R | 000087F |
| AGAP012438 | 2R | 000088F |
| AGAP012488 | 2R | 000088F |
| AGAP028987 | 2R | 000088F |
| AGAP012442 | 2L | 000089F |
| AGAP012441 | 2L | 000089F |
| AGAP012760 | 2L | 000089F |
| AGAP012440 | 2L | 000089F |
| AGAP029002 | 2L | 000093F |
| AGAP029003 | 2L | 000093F |
| AGAP012782 | 2L | 000093F |
| AGAP012784 | 2L | 000093F |
| AGAP012625 | 2L | 000093F |

| | | |
|---|---|---|
| AGAP012515 | 2L | 000093F |
| AGAP012516 | 2L | 000093F |
| AGAP012517 | 2L | 000093F |
| AGAP012624 | 2L | 000093F |
| AGAP012876 | 3L | 000095F |
| AGAP012663 | 3L | 000095F |
| AGAP012891 | 2L | 000096F |
| AGAP012449 | 2R | 000104F |
| AGAP012484 | 2L | 000105F |
| AGAP012538 | 2L | 000105F |
| AGAP012506 | 2L | 000105F |
| AGAP012481 | 2R | 000106F |
| AGAP012610 | 2R | 000106F |
| AGAP012912 | 2R | 000106F |
| AGAP028997 | X | 000107F |
| AGAP028838 | X | 000107F |
| AGAP028835 | X | 000107F |

| | | |
|---|---|---|
| AGAP028832 | X | 000107F |
| AGAP028830 | X | 000107F |
| AGAP028831 | X | 000107F |
| AGAP028992 | X | 000107F |
| AGAP028850 | X | 000107F |
| AGAP028851 | X | 000107F |
| AGAP028852 | X | 000107F |
| AGAP028854 | X | 000107F |
| AGAP028857 | X | 000107F |
| AGAP028872 | X | 000107F |
| AGAP028873 | X | 000107F |
| AGAP028876 | X | 000107F |
| AGAP028877 | X | 000107F |
| AGAP028874 | X | 000107F |
| AGAP028875 | X | 000107F |
| AGAP028878 | X | 000107F |
| AGAP028879 | X | 000107F |

| | | |
|---|---|---|
| AGAP028836 | X | 000107F |
| AGAP028935 | X | 000107F |
| AGAP028934 | X | 000107F |
| AGAP028932 | X | 000107F |
| AGAP028930 | X | 000107F |
| AGAP028938 | X | 000107F |
| AGAP028911 | X | 000107F |
| AGAP028917 | X | 000107F |
| AGAP028954 | X | 000107F |
| AGAP028957 | X | 000107F |
| AGAP028882 | X | 000107F |
| AGAP028880 | X | 000107F |
| AGAP028984 | X | 000107F |
| AGAP028945 | X | 000107F |
| AGAP028921 | X | 000107F |
| AGAP028788 | X | 000107F |
| AGAP028789 | X | 000107F |

| | | |
|---|---|---|
| AGAP028787 | X | 000107F |
| AGAP028802 | X | 000107F |
| AGAP028809 | X | 000107F |
| AGAP028808 | X | 000107F |
| AGAP028827 | X | 000107F |
| AGAP028959 | X | 000107F |
| AGAP028958 | X | 000107F |
| AGAP029034 | X | 000107F |
| AGAP029037 | X | 000107F |
| AGAP028843 | X | 000107F |
| AGAP028841 | X | 000107F |
| AGAP028840 | X | 000107F |
| AGAP028847 | X | 000107F |
| AGAP028845 | X | 000107F |
| AGAP028844 | X | 000107F |
| AGAP028988 | X | 000107F |
| AGAP028871 | X | 000107F |

| | | |
|---|---|---|
| AGAP028866 | X | 000107F |
| AGAP028860 | X | 000107F |
| AGAP028869 | X | 000107F |
| AGAP028949 | X | 000107F |
| AGAP028925 | X | 000107F |
| AGAP028926 | X | 000107F |
| AGAP028920 | X | 000107F |
| AGAP028923 | X | 000107F |
| AGAP028906 | X | 000107F |
| AGAP028907 | X | 000107F |
| AGAP028902 | X | 000107F |
| AGAP028961 | X | 000107F |
| AGAP028965 | X | 000107F |
| AGAP028942 | X | 000107F |
| AGAP028940 | X | 000107F |
| AGAP028941 | X | 000107F |
| AGAP028946 | X | 000107F |

| | | |
|---|---|---|
| AGAP028910 | X | 000107F |
| AGAP028916 | X | 000107F |
| AGAP028918 | X | 000107F |
| AGAP028898 | X | 000107F |
| AGAP028899 | X | 000107F |
| AGAP028894 | X | 000107F |
| AGAP028890 | X | 000107F |
| AGAP028891 | X | 000107F |
| AGAP028893 | X | 000107F |
| AGAP028798 | X | 000107F |
| AGAP028796 | X | 000107F |
| AGAP028950 | X | 000107F |
| AGAP028939 | X | 000107F |
| AGAP028884 | X | 000107F |
| AGAP028812 | X | 000107F |
| AGAP012444 | 3R | 000111F |
| AGAP012445 | 3R | 000111F |

| | | |
|---|---|---|
| AGAP012443 | 3R | 000111F |
| AGAP028575 | 3R | 000111F |
| AGAP028579 | 3R | 000119F |
| AGAP028931 | 2L | 000120F |
| AGAP029033 | 2L | 000120F |
| AGAP028989 | 2L | 000120F |
| AGAP012776 | 2L | 000124F |
| AGAP012766 | 3R | 000128F |
| AGAP012762 | 2L | 000133F |
| AGAP012588 | 3R | 000142F |
| AGAP012494 | 2R | 000145F |
| AGAP012532 | 2R | 000145F |
| AGAP012533 | 2R | 000154F |
| AGAP012500 | 2R | 000155F |
| AGAP012499 | 2R | 000155F |
| AGAP012583 | 2R | 000206F |
| AGAP012526 | 2R | 000206F |

| | | |
|---|---|---|
| AGAP028915 | UNKN | 000243F |
| AGAP028914 | UNKN | 000243F |
| AGAP028824 | X | 000248F |
| AGAP028955 | X | 000248F |
| AGAP028887 | X | 000248F |
| AGAP028886 | X | 000248F |
| AGAP028881 | X | 000248F |
| AGAP012700 | 2L | 000283F |
| AGAP012752 | 2L | 000283F |
| AGAP012548 | not placed | N/A |
| AGAP012738 | not placed | N/A |
| AGAP012727 | not placed | N/A |
| AGAP028913 | not placed | N/A |
| AGAP012507 | not placed | N/A |
| AGAP029100 | not placed | N/A |
| AGAP012751 | not placed | N/A |
| AGAP012874 | not placed | N/A |

| | | |
|---|---|---|
| AGAP012778 | not placed | N/A |
| AGAP012815 | not placed | N/A |
| AGAP028582 | not placed | N/A |
| AGAP012468 | not placed | N/A |
| AGAP012467 | not placed | N/A |
| AGAP012461 | not placed | N/A |
| AGAP012672 | not placed | N/A |
| AGAP012460 | not placed | N/A |
| AGAP028268 | not placed | N/A |
| AGAP012448 | not placed | N/A |
| AGAP012897 | not placed | N/A |
| AGAP028295 | not placed | N/A |
| AGAP028888 | not placed | N/A |
| AGAP012466 | not placed | N/A |
| AGAP028786 | not placed | N/A |
| AGAP012455 | not placed | N/A |
| AGAP012551 | not placed | N/A |

| | | |
|---|---|---|
| AGAP012446 | not placed | N/A |
| AGAP012909 | not placed | N/A |
| AGAP028801 | not placed | N/A |
| AGAP012923 | not placed | N/A |
| AGAP012502 | not placed | N/A |
| AGAP012940 | not placed | N/A |
| AGAP012615 | not placed | N/A |
| AGAP012701 | not placed | N/A |
| AGAP028865 | not placed | N/A |
| AGAP028650 | not placed | N/A |
| AGAP013132 | not placed | N/A |
| AGAP012485 | not placed | N/A |
| AGAP012539 | not placed | N/A |
| AGAP012744 | not placed | N/A |
| AGAP012483 | not placed | N/A |
| AGAP028908 | not placed | N/A |
| AGAP028904 | not placed | N/A |

| | | |
|---|---|---|
| AGAP028905 | not placed | N/A |
| AGAP028901 | not placed | N/A |
| AGAP012733 | not placed | N/A |
| AGAP012862 | not placed | N/A |
| AGAP028574 | not placed | N/A |
| AGAP012452 | not placed | N/A |
| AGAP028960 | not placed | N/A |
| AGAP028964 | not placed | N/A |
| AGAP012592 | not placed | N/A |
| AGAP012597 | not placed | N/A |
| AGAP012848 | not placed | N/A |
| AGAP012840 | not placed | N/A |
| AGAP012846 | not placed | N/A |
| AGAP012601 | not placed | N/A |
| AGAP012621 | not placed | N/A |
| AGAP029036 | not placed | N/A |
| AGAP029038 | not placed | N/A |

| | | |
|---|---|---|
| AGAP012889 | not placed | N/A |
| AGAP028246 | not placed | N/A |
| AGAP028791 | not placed | N/A |
| AGAP028795 | not placed | N/A |
| AGAP012884 | not placed | N/A |
| AGAP012463 | not placed | N/A |
| AGAP012794 | not placed | N/A |
| AGAP012916 | not placed | N/A |
| AGAP012935 | not placed | N/A |

avro==1.8.2
certifi==2018.4.16
chardet==3.0.4
ConsensusCore==1.0.2
ConsensusCore2==3.0.0
Cython==0.28.5
decorator==4.3.0
edlib==1.2.3.post1
falcon-kit==1.2.2+git.00e8272b663d32a0962ae92ab92324a3b3eb4b46
FALCON-pbsmrtpipe==1.0.0
FALCON-polish==1.0.0
falcon-unzip==1.1.2+git.354e53c5beb103c5c8301914d86fe96364b29288
functools32==3.2.3.post2
future==0.16.0
GenomicConsensus==2.2.2
idna==2.7
intervaltree==2.1.0
iso8601==0.1.12
jsonschema==2.6.0
msgpack==0.5.6
msgpack-python==0.5.6
networkx==2.1
numpy==1.15.0
pbalign==0.3.0
pbcommand==1.1.2
pbcore==1.5.4
pbcoretools==0.2.4
pypeflow==2.0.4+git.005acb16689c18c09cf552b42911e69629ffeceb
pysam==0.13
pytz==2018.5
requests==2.19.1
sortedcontainers==2.0.4
urllib3==1.23
xmlbuilder==1.0

Note S1.
Software versions for *de novo* assembly with FALCON-Unzip.

```
[job.defaults]
njobs = 100
submit = qsub -S /bin/bash -sync y -V  \
  -q ${JOB_QUEUE}     \
  -N ${JOB_NAME}        \
  -o "${JOB_STDOUT}" \
  -e "${JOB_STDERR}" \
  -pe smp ${NPROC}    \
  "${JOB_SCRIPT}"

JOB_QUEUE = default7
MB = 30000
NPROC = 6
[General]
input_type = raw
input_fofn = fasta.fofn
pwatcher_type=blocking

genome_size = 280000000
seed_coverage = 40
length_cutoff = -1
length_cutoff_pr = 5000
falcon_greedy = False
falcon_sense_greedy=False

pa_daligner_option = -e0.76 -l1200 -k18 -h70 -w8 -s100
ovlp_daligner_option = -k24 -h1024 -e.95 -l1800 -s100
pa_HPCdaligner_option = -v -B128 -M24
ovlp_HPCdaligner_option = -v -B128 -M24
pa_HPCTANmask_option = -k18 -h480 -w8 -e.8 -s100
pa_HPCREPmask_option = -k18 -h480 -w8 -e.8 -s100
pa_DBsplit_option = -x500 -s400
ovlp_DBsplit_option = -s400

falcon_sense_option = --output-multi --min-idt 0.70 --min-cov 3 --max-n-read 400 --n-core 24
overlap_filtering_setting = --max-diff 100 --max-cov 150 --min-cov 3 --n-core 24

[job.step.da]
NPROC=4
MB=20000
njobs=100
[job.step.la]
NPROC=8
MB=40000
njobs=100
[job.step.cns]
NPROC=8
MB=40000
njobs=100
```

```
[job.step.pda]
NPROC=8
MB=40000
njobs=100
[job.step.pla]
NPROC=4
MB=20000
njobs=100
[job.step.asm]
NPROC=24
MB=120000
```

Note S2.
Configuration file for FALCON assembly.

```
[General]

[Unzip]
input_fofn= fasta.fofn
input_bam_fofn=bam.fofn

[job.defaults]
pwatcher_type=blocking
max_n_open_files = 1000
njobs=50
JOB_QUEUE=bigmem
NPROC=4
MB=32000
submit = qsub -S /bin/bash -sync y -V  \
  -q ${JOB_QUEUE}     \
  -N ${JOB_NAME}        \
  -o "${JOB_STDOUT}" \
  -e "${JOB_STDERR}" \
  -pe smp ${NPROC}    \
  "${JOB_SCRIPT}"
[job.step.unzip.track_reads]
njobs=1
NPROC=24
MB=192000
[job.step.unzip.blasr_aln]
njobs=45
NPROC=24
MB=192000
[job.step.unzip.phasing]
njobs=100
NPROC=2
MB=16000
[job.step.unzip.hasm]
njobs=1
NPROC=48
MB=384000
[job.step.unzip.quiver]
njobs=45
NPROC=24
MB=192000
```

### Note S3.
### Configuration file for FALCON-Unzip.
